# The Dual Action of Human Antibodies Specific to *Plasmodium falciparum* PfRH5 and PfCyRPA: Blocking Invasion and Inactivating Extracellular Merozoites

**DOI:** 10.1101/2023.02.07.527440

**Authors:** Greta E. Weiss, Robert J. Ragotte, Doris Quinkert, Amelia M. Lias, Madeline G. Dans, Coralie Boulet, Barnabas G. Williams, Brendan S. Crabb, Simon J. Draper, Paul R. Gilson

## Abstract

The *Plasmodium falciparum* reticulocyte-binding protein homolog 5 (PfRH5) is the current leading blood-stage malaria vaccine candidate. PfRH5 functions as part of the pentameric PCRCR complex containing PTRAMP, CSS, PfCyRPA and PfRIPR, all of which are essential for infection of human red blood cells (RBCs). To trigger RBC invasion, PfRH5 engages with RBC protein basigin in a step termed the RH5-basigin binding stage. Although we know increasingly more about how antibodies specific for PfRH5 can block invasion, much less is known about how antibodies recognizing other members of the PCRCR complex can inhibit invasion. To address this, we performed live cell imaging using monoclonal antibodies (mAbs) which bind PfRH5 and PfCyRPA. We measured the degree and timing of the invasion inhibition, the stage at which it occurred, as well as subsequent events. We show that parasite invasion is blocked by individual mAbs, and the degree of inhibition is enhanced when combining a mAb specific for PfRH5 with one binding PfCyRPA. In addition to directly establishing the invasion-blocking capacity of the mAbs, we identified a secondary action of certain mAbs on extracellular parasites that had not yet invaded where the mAbs appeared to inactivate the parasites by triggering a developmental pathway normally only seen after successful invasion. These findings suggest that epitopes within the PfCyRPA-PfRH5 sub-complex that elicit these dual responses may be more effective immunogens than neighboring epitopes by both blocking parasites from invading and rapidly inactivating extracellular parasites. These two protective mechanisms, prevention of invasion and inactivation of uninvaded parasites, resulting from antibody to a single epitope indicate a possible route to the development of more effective vaccines.

**Author Summary:** Malaria is a sometimes-fatal disease caused by protozoan parasites of which *Plasmodium falciparum* is the most deadly species that causes hundreds of millions of infections and half a million deaths per year. A partially effective vaccine is available to block parasite forms transmitted by mosquitoes but not the subsequent blood stage which causes symptomatic disease. To fight blood stage parasites, proteins have been identified such as PfRH5, that aid parasite entry into human red blood cells (RBCs) and vaccines made from these proteins can trigger the production of antibodies that bind the parasite proteins thereby blocking RBC invasion. PfRH5 forms a complex with another parasite protein called PfCyRPA and together antibodies to PfCyRPA and PfRH5 are highly effective in reducing parasite growth. Here we investigated how antibodies to PfCyRPA and PfRH5 actually block invasion using video microscopy of live parasites. As anticipated, we found the antibodies not only stopped most parasites from invading but of those parasites that did invade, they took longer to do so, suggesting the antibodies were physically inhibiting the invasion process. One unanticipated effect of both PfRH5 antibodies and one of three PfCyRPA antibodies tested, was that they triggered the uninvaded parasites to change into cellular forms normally only seen inside RBCs. These intracellular forms are no longer competent to invade and so the PfRH5/CyRPA antibodies have the potential of both neutralize parasites by physically preventing RBC entry and by changing the parasites into invasion incompetent forms.

## Introduction

Globally, malaria remains a serious problem with 247 million cases of malaria worldwide, and 619,000 deaths in 2021 [1], with the majority of disease burden and death due to *Plasmodium falciparum* (*Pf*). Despite a growing arsenal of antimalarial drugs, it is unlikely that drugs alone will be enough to eradicate malaria. Disruptions to pharmaceutical supply chains caused an increase in malaria disease burden during the COVID-19 pandemic, further underlining the importance of a highly effective malaria vaccine, which would continue to provide protection when medicines are not available, or when a constant medical presence is not possible.

Antibodies are known to play a key role in protection from malaria [2]. Low-level antibody responses to parasite invasion protein PfRH5, can be naturally-acquired following many years of malaria exposure and associate with clinical immunity and inhibit parasite growth *in-vitro* [3]. High levels of PfRH5 vaccine-induced antibodies can block merozoite invasion of erythrocytes [4], and are associated with delayed time to diagnosis in a controlled human malaria infection trial in vaccinated individuals [5]. PfRH5 forms a chain-like PCRCR complex with *P. falciparum* Cysteine-Rich Protective Antigen (PfCyRPA), PfRH5-interacting protein (PfRIPR), *P. falciparum* cysteine-rich small secreted protein (PfCSS) and *P. falciparum* Plasmodium thrombospondin-related apical merozoite protein (PfPTRAMP) [6–8], which has been shown to localize at the tight junction between the merozoite and erythrocyte immediately prior to invasion [6, 9]. PfRH5, PfCyRPA, PfRIPR, PfCSS and PfPTRAMP are required for merozoite invasion of erythrocytes with PfRH5, PfCyRPA, PfRIPR each able to stimulate the production of cross-strain neutralizing antibodies [6, 9–15]. Of the five PCRCR components, PfRH5, PfCyRPA and PfRIPR are the most well characterized and are compelling malaria vaccine candidates with experimental vaccination of non-human primates and humans with PfRH5 producing potent neutralizing antibodies [16, 17]. Understanding the effect of antibodies targeting the PCRCR complex antigens, both in isolation and combination, is now critical to effective next-generation blood-stage vaccine design. Importantly, the amount of antibody required for protection may be greatly reduced by exploiting synergistic interactions between antibodies.

Antibodies recognizing the same complex could be expected to synergize through a wide variety of mechanisms, including, but not limited to, causing a conformational change that enhances binding of a second antibody [18] slowing down invasive processes allowing other antibodies more time to bind [4] or by stabilizing antibody binding through lateral interactions [19]. The ideal effect of antibodies would be rapid and lead to permanent disabling of the parasites’ ability to invade or survive. While antibody binding to a parasite might temporarily prevent invasion, if that parasite is not disabled by this interaction, then the dissociation of that antibody could leave the parasite virulent and able to attempt invasion again.

In this study we use live cell imaging to examine the visible temporal and morphological effects of mAbs binding to different epitopes of PfRH5 and PfCyRPA on invasion, the post-invasion development of young parasites, and the inactivation of uninvaded merozoites. While the ability of PCRCR antigen-specific antibodies to block invasion is known, this is the first report of their capacity to inactivate uninvaded parasites. This work illuminates new mechanisms through which neutralizing antibodies to certain epitopes of PfRH5 and PfCyRPA function and future studies of the cellular and molecular basis of these observations could open the door to strategies for the rational design of a highly effective blood-stage malaria vaccine.

## Results

### Antibody inhibition of merozoite invasion

To characterize the effects of mAbs specific for PfRH5 and PfCyRPA on invasion of erythrocytes, live cell microscopy was performed on clonal 3D7 parasites using methods that had been previously developed [20]. As expected, parasites in the presence of the control EBL 040 mAb (targeting *Ebola virus*)[4, 21] followed the usual progression of invasion with distinct stages marked first by egress of merozoites from the schizont (Fig 1A, S1 Table, S1 Video) [20, 22, 23], contact of the merozoite with the erythrocyte, and deformation of the erythrocyte membrane. Following this is the short stage where the PCRCR complex acts and the tight junction forms between the merozoite and erythrocyte. Here we term this the “RH5-basigin binding stage”, and merozoite invasion of the erythrocyte ensues over the next 10 seconds. After this, temporary changes to the invaded erythrocyte occur called echinocytosis, and the merozoite differentiates into an intracellular ring (Fig 1A, S1 Video).

**Fig 1.**
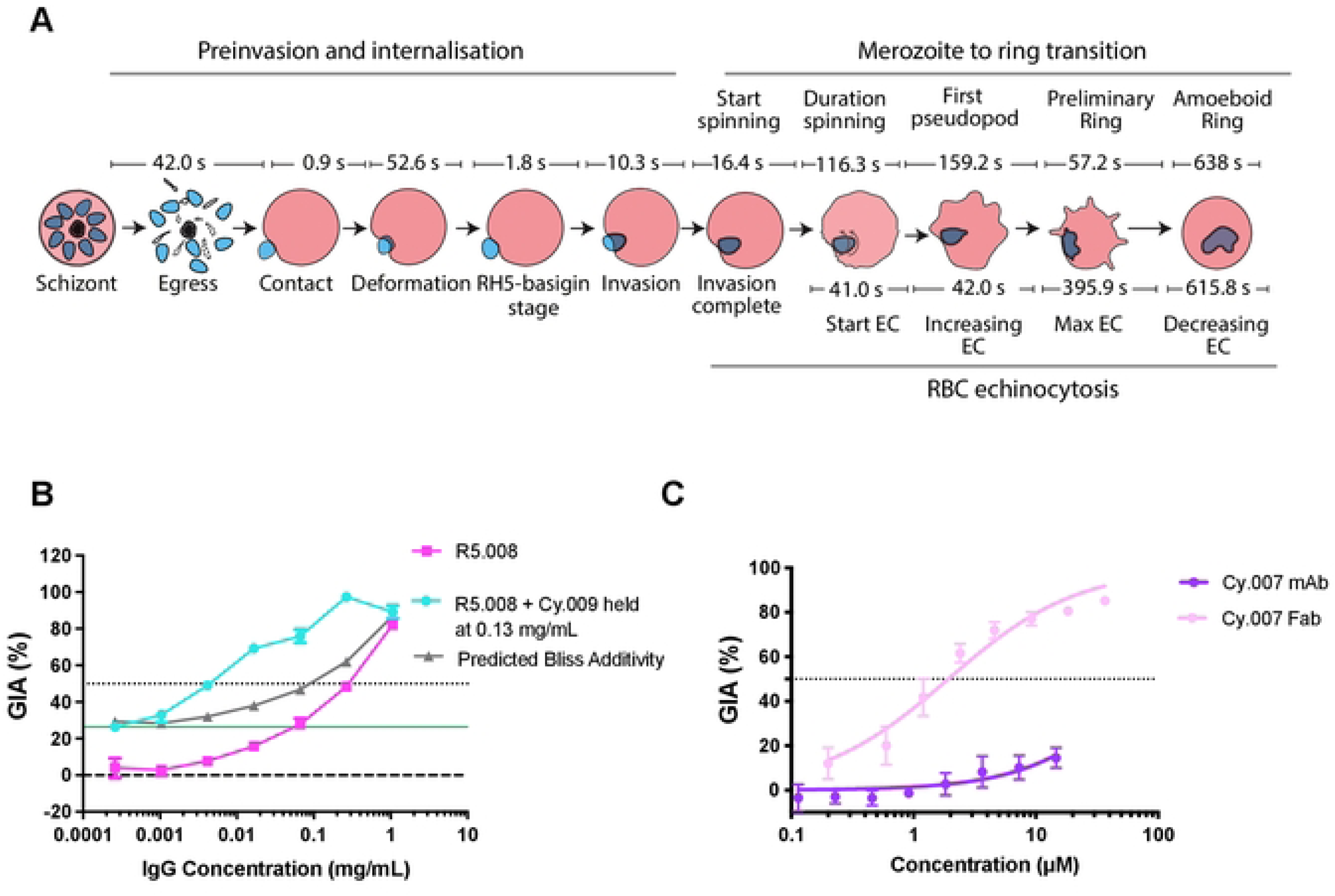
The major stages of invasion of *Plasmodium falciparum* blood stage parasites and growth inhibitory activity (GIA) of anti-PfRH5 and -PfCyRPA IgGs and Fab fragments. (A) Several videos of *P. falciparum* in the presence of 400 µg/mL EBL 040 control IgG were analyzed to derive the average times for each of the major steps of pre-invasion, internalization, merozoite to ring transition and echinocytosis (EC) of the infected erythrocyte. After egress, merozoites took 42.0 s to contact the erythrocyte they invaded (all times are means). Following an initial contact of 0.9 s, the merozoites deformed their erythrocytes for 52.6 s followed by the quiescent complex stage of 1.8 s, when it is believed the PCRCR complex and tight junction form (“Rh5-basigin stage”). Merozoite internalization or invasion takes 10.3 s after which the merozoite starts to spin 16.4 s later. Spinning lasts 116.3 s and once finished, a pseudopod emerges from the merozoite 159.2 s later. 57.2 s after this, the merozoite membrane starts to become irregular and by 638 s a fully formed amoeboid ring is evident. 41 s after invasion is complete, the host erythrocyte develops membranous protrusions in a process called echinocytosis. These protrusions increase in their extent reaching a maximum after 42 s. This state continues for 395.9 s until the erythrocyte returns to its usual biconcave shape. Of note, the timing and duration of echinocytosis is highly variable. (B) *In vitro* single cycle growth inhibition activity (GIA) assay dilution series of R5.008 mAb alone (pink), or in combination with Cy.009 mAb held at 0.13 mg/mL (blue), against 3D7 parasites. Predicted Bliss additivity is indicated (grey). The solid green line indicates the GIA of mAb Cy.009 held alone at a fixed concentration of 0.13 mg/mL. Dotted line indicates 50% GIA. (C) Comparison of the GIA of the Cy.007 mAb (purple) with its Fab fragment (light pink) indicating the latter is much more potent. Assay performed as in B.

Anti-PfRH5 mAbs were isolated from volunteers immunized in the first human Phase Ia PfRH5 vaccine trial [24]. Individual mAbs were characterized: clone R5.004 was found to be strongly neutralizing and to bind directly to the basigin-binding site of PfRH5 [4]; whilst clone R5.008 was found to be moderately neutralizing possibly by hindering PfRH5’s access to basigin through steric clashes with the RBC membrane or basigin’s RBC binding partners PCMA or MCT1 (S1 Fig A and B) [4, 25]. It has recently been discovered that PfRH5 must have its pro-sequence cleaved by plasmepsin X before it can engage basigin [26].

We also analyzed three mAbs which bind PfCyRPA and were composed of variable regions from vaccinated chickens and a human IgG1 constant region [19]. These three mAbs all bind to the beta propeller blade 1 and 2 regions of PfCyRPA (S1 Fig C and D). PfCyRPA-binding mAbs demonstrated modest levels of inhibition [7]. However, in growth inhibition activity (GIA) assays, R5.008 and Cy.009 synergized to strongly inhibit invasion (Fig 1B). Because of this, R5.008 was filmed at a concentration causing less inhibition to directly compare results with the R5.008 + Cy.009 combination, while other individual mAbs were filmed at concentrations resulting in an intermediate level of inhibition to assess the stages of invasion that were affected. We also evaluated PfCyRPA fragment antigen binding (Fab) molecules in GIA assays and found the Fabs consistently demonstrated greater GIA than whole IgGs at equivalent molarities (Fig 1C, S1 Fig E-G). Since this was especially evident for Cy.007, with Fabs being about >10-fold more inhibitory than whole IgGs, we included this Fab in our live imaging analysis.

Videos of egress and invasion were analyzed in detail for each of the mAbs, Fabs and the control EBL 040 antibody with a median of 11 videos analyzed for each antibody. The concentration used are indicated in brackets (in µg/mL). The number of merozoite-erythrocyte contacts per egress was noted and was equal to or greater than that in the control antibody, indicating the parasites were healthy (Fig 2A, S2A Table). In addition, the times from egress to first merozoite contact with erythrocytes were similar demonstrating the experimental conditions were consistent for each antibody combination (Fig S2A, S5A Table). With the exception of R5.004-(22) the antibodies did not greatly reduce the degree with which the merozoites deformed their target RBCs which is thought to be a product of merozoite ligands acting upstream of the PCRCR complex and the actomyosin invasion motor (S2B Fig, S5B Table) [20].

**Fig 2.**
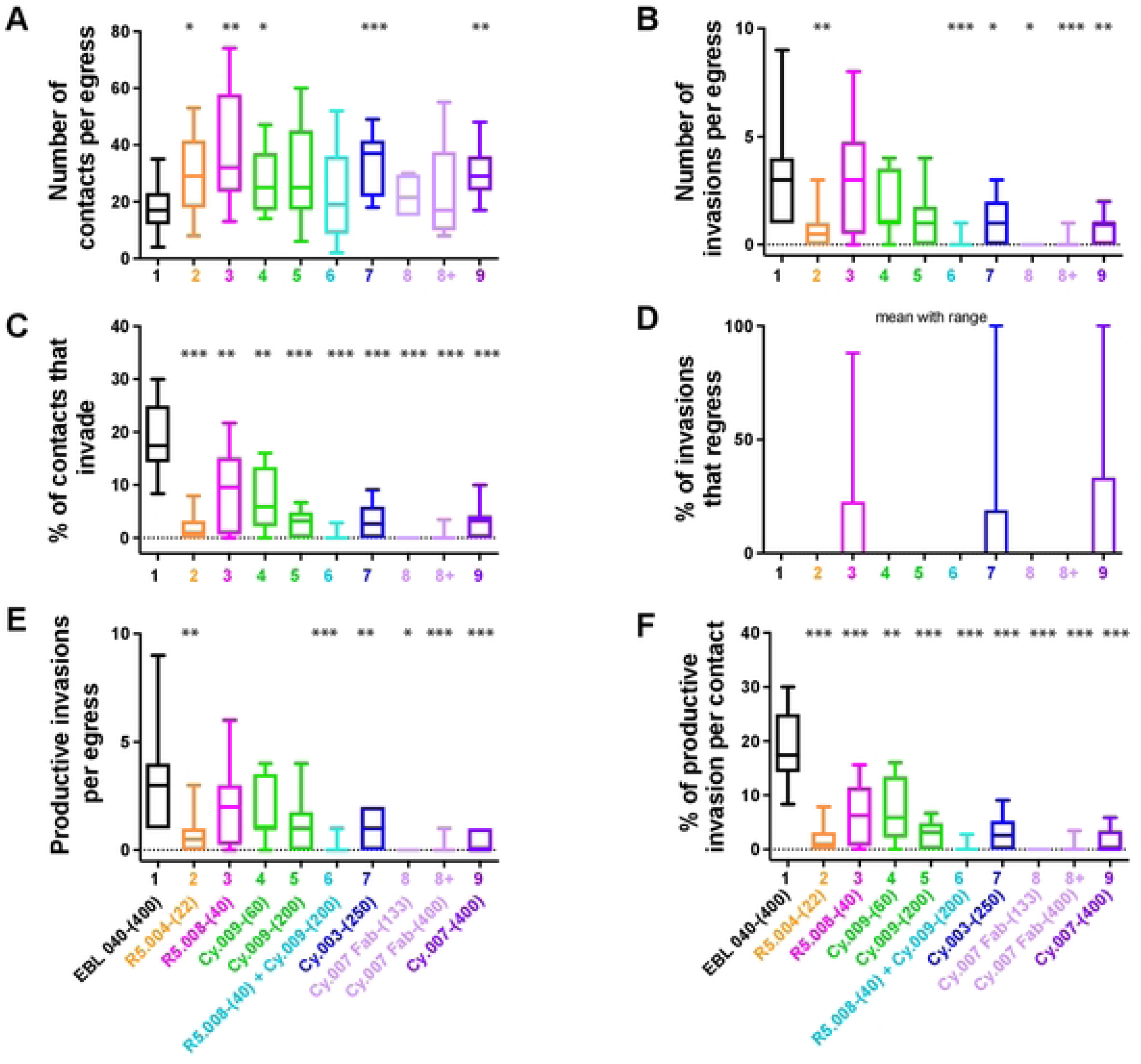
Parasite specific mAbs to PfRH5 and PfCyRPA inhibit *Plasmodium falciparum* invasion of human erythrocytes. (A-F) Several live cell videos of *P. falciparum* merozoites egressing and attempting to invade erythrocytes in the presence of each of the antibody (concentrations and combinations indicated) were analyzed. The number of successful events is presented for each parameter indicated by the y axis. Full antibody names and concentrations (µg/mL) are indicated below bottom graphs. The boxes indicate the median number of events (A,B,E) or the percentage of events (C,D,F) and the 25% to 75% percentile, and whiskers show the range. Statistical analyses were performed using unpaired t tests in GraphPad PRISM V 9.0. The asterisks indicate where parasite mAbs have altered the number/percentage of events significantly from the EBL 040 control with *p<0.05, **p<0.01 and ***p<0.001.

The number of invasions per egress was reduced in the R5.004-(22), Cy.003-(250) and Cy.007 antibodies as well as in the R5.008-(40) + Cy.009-(200) combination as compared to the control EBL 040-(400) (Fig 2B, S2B Table, S1-S10 Videos). As the direction of schizont egress and the number of erythrocytes available for invasion can vary, a complementary measure of invasion was also used: the percentage of merozoites that invade after contacting an erythrocyte. Using this measure, invasion was reduced for all anti-PfRH5 and -PfCyRPA antibodies compared to the control (Fig 2C, S2C Table). R5.004-(22), Cy.007 Fab-(133) and -(400), and the combination of R5.008-(40) + Cy.009-(200) were so inhibitory, at 97.5-100% inhibition, that these yielded limited data (Fig 2C). In three experimental conditions, R5.008-(40), Cy.003-(250) and Cy.007-(400), some merozoites regressed back out of the erythrocyte a short time after invasion, which we have referred to as “regression” (Fig 2D, S2D Table). To account for this we assessed “productive invasions” which are those invasions where the merozoite remains inside the erythrocyte during the 20-minute observation period (Fig 2E). Overall, the number of productive invasions per egress and the percentage of productive invasions per contact showed similar trends to the total numbers of invasions per egress and percentage of invasions per contact (Fig 2B versus 2E and Fig 2C versus 2F, S2E-F Table).

Importantly, for R5.004(22), Cy.009(200), Cy.003(250) and Cy.007(400) there were no statistically significant differences when comparing the number of invasions per egress, productive invasions per egress, or the percent of contacts which invaded or resulted in productive invasions (Fig 2, S2 Tables). This allowed us to proceed comparing the different mechanisms of invasion inhibition which occur with these various mAbs, given the overall level of invasion inhibition is comparable whilst the invasion stages affected by the mAbs could vary.

### Antibody effects on early pre-invasion interactions

Pre-invasion consists of the time from first contact between the erythrocyte and merozoite to the start of invasion, with early pre-invasion events including initial contact and erythrocyte deformation (Fig 1A). The antibodies did not greatly change the short period of time from first merozoite contact to the start of erythrocyte deformation, although R5.004-(22) was slightly shortened relative to Cy.009-(200) and Cy.007-(400) (Fig 3A, S3A Table). R5.004-(22) also shortened the duration of deformation (mean 8.3 s) compared to PfCyRPA-binding mAbs (Fig 3B, S3B Table). Although Cy.007-(400) prolonged deformation (mean: 107.9 s) compared to R5.004- (22), R5.008-(40) and Cy.009-(200), this was not significantly different to EBL 040 control (mean: 52.7 s) (Fig 3B, S3B Table). The CyRPA-binding mAb Cy.007-(400) prolonged the whole pre-invasion period from first contact to the start of invasion in comparison to the control antibody, RH5-binding mAbs and Cy.009-(60) and Cy.003-(250) (Fig 3C, S3C Table).

**Fig 3.**
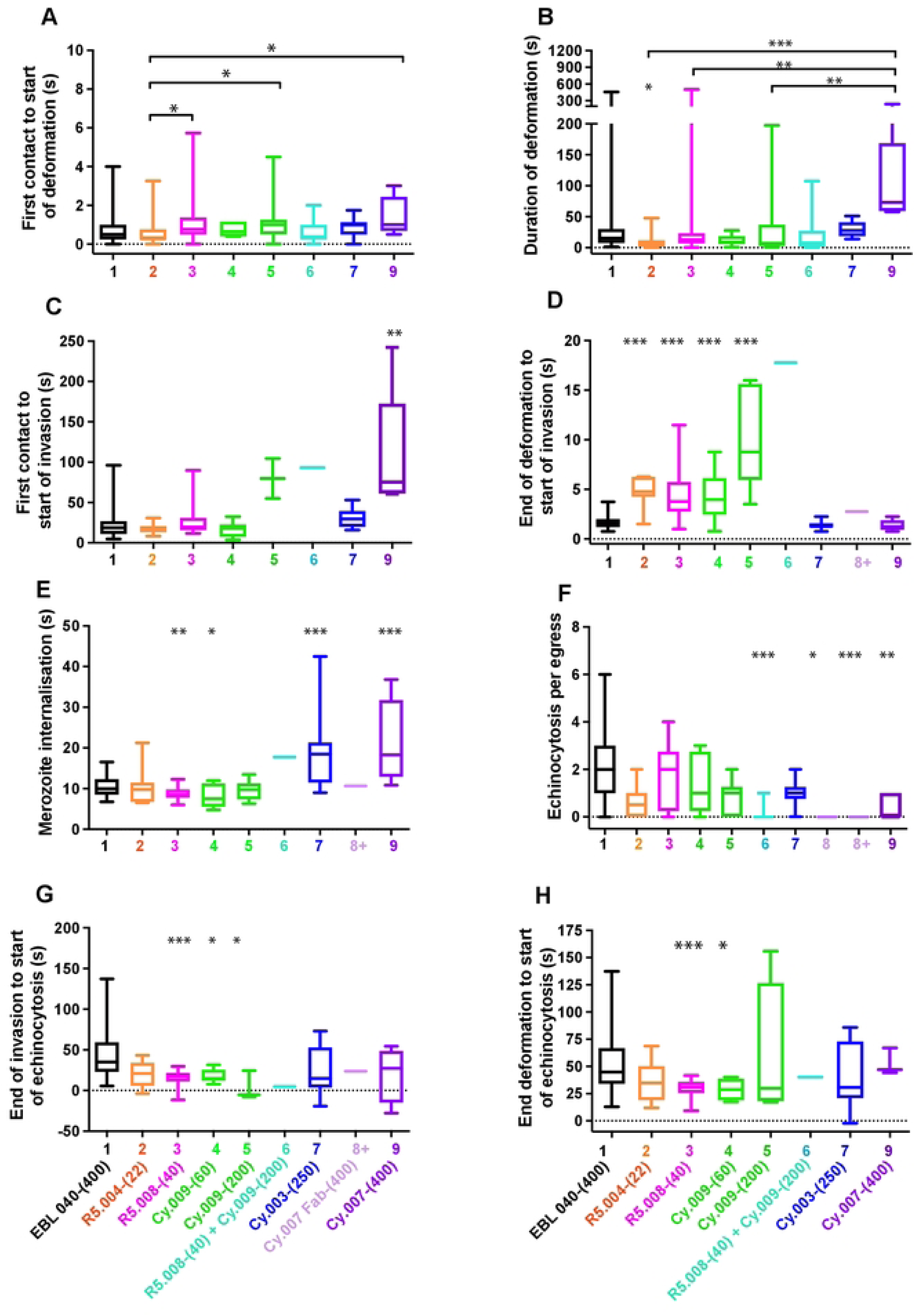
Antibodies to PfRH5 and PfCyRPA modify temporal aspects of the pre- and post- invasion phases of *Plasmodium falciparum* into human erythrocytes. (A-E) The invasion steps being monitored from live cell videos of antibody-treated parasites are indicated on the y-axes. The antibody types and concentrations are indicated on the x-axes. Anti-PfRH5 (orange, pink) and Cy.009 (green) antibodies tend to increase the length of the preinvasion phase from first erythrocyte contact to the start of merozoite penetration. The Cy.003 (dark blue) and Cy.007 IgG and Fabs (purples) tend to increase the length of the merozoite internalization period. (F) Cy.007 antibodies decrease the numbers of echinocytosis events per egress and (G-H) R5.008 (pink) and Cy.009 (green) decrease the time from the end of invasion/deformation to echinocytosis. Full antibody names and concentrations (µg/mL) are indicated below bottom graphs. Antibody 8 indicates Cy.007 Fab-(133). The boxes indicate the median number of events and the 25% to 75% percentile, and whiskers show the range. Statistical analyses were performed using unpaired t tests in GraphPad PRISM V 9.0. The asterisks indicate where parasite mAbs have altered the number of events significantly from the EBL 040 control with *p<0.05, **p<0.01 and ***p<0.001.

### Antibody effects on the complex stage; late pre-invasion interactions

The RH5-basigin binding stage, the time from the end of deformation to the start of invasion, is the time PCRCR is thought to have its primary function, aiding rhoptry release and tight junction formation [6, 9, 20]. This stage of invasion is tightly regulated, with an average time of 1.8 to 2.0 s (Fig 1A, S1 Table) [20]. In comparison to control antibody EBL 040, both R5.004-(22) and R5.008- (40), as well as Cy.009-(60 and 200) caused a delay at this stage, that was dose dependent in the case of Cy.009 (Fig 3D, S3D Table). Although there was only a single instance of an invasion under these conditions, the R5.008-(40) + Cy.009-(200) combination resulted in the longest delay, which at 17.75 s was slightly longer than the sum of the means of each antibody alone (14.27 s). In contrast, the PfCyRPA-binding Cy.003-(250) and Cy.007 antibodies did not cause a delay at this stage of invasion (Fig 3D).

### Antibody effects on invasion

Internalization of the parasite, measured from the time the merozoite begins to enter the erythrocyte until it is entirely inside the erythrocyte, typically has little variability and consistently takes about 10 s (Fig 1A, S1 Table) [20]. The Cy.003-(250) and Cy.007-(400) significantly slowed internalization, causing this stage to take approximately double the time observed in the presence of control antibody or the other individual antibodies (Fig 3E, S3E Table). Although R5.008-(40) and Cy.009-(60) decreased the time of internalization, the combination of these two antibodies greatly slowed the invasion time, although this combination was so inhibitory there was only a single invasion observed (Fig 3E). When the RH5-basigin binding stage and invasion stages were combined there was an overall increase in the time from the end of deformation to the end of invasion for R5.004-(22), Cy.009-(200), Cy.003-(250) and Cy.007-(400) compared to the control (S2C Fig, S5C Table).

### Antibody effects on echinocytosis

After invasion is completed, the invaded erythrocyte typically becomes spherical with distinctive spikes covering its surface for several minutes before returning to its usual biconcave shape [22, 23] (Fig 1A). This process is called echinocytosis and is strongly associated with successful invasions [20]. We noted that there were significant reductions in the numbers of these events per egress for the R5.008 and Cy.009 combination as well as the Cy.007 Fabs and IgGs (Fig 3F and S3F Table) like the invasions per egress data (Fig 2B). The echinocytosis stage of the invasion process has wide natural variability spanning several minutes. For every individual mAb, there were instances of echinocytosis starting before invasion had completed, something rarely observed previously [20, 27] (negative values in Fig 3G and S3G Table). Echinocytosis was triggered earlier for parasites in the presence of R5.008-(40) (mean 15.0 s) and Cy.009-(60 and 200) (mean 18.00 and 3.8 s, respectively) compared to EBL 040 (mean 40.9 s)(Fig 3G and S3G Table). Since PfRH5 likely binds basigin at the end of deformation [20], and PCRCR-complex binding has been linked to echinocytosis [27], we examined this time period (end of deformation to start of echinocytosis). R5.008-(40) and Cy.009-(60) caused a decrease in this time (Fig 3H and S3H Table). Although the CyRPA antibodies tended to increase the mean duration of echinocytosis this was only statistically significant for Cy.003-(250) (S2D Fig and S5D Table).

To further analyze echinocytosis of the newly invaded RBCs, we measured increasing echinocytosis, the time from the initiation of echinocytosis to the point when maximum echinocytosis was reached, the duration of maximum echinocytosis, and decreasing echinocytosis, the time it took for the erythrocyte to recover its usual biconcave shape (Fig 1A, S1 Table). For the EBL 040 control, increasing echinocytosis lasted a mean of 42.0 s, maximum echinocytosis 395.9 s, and decreasing echinocytosis 615.8 s (S2E-G Fig, S1 Table, S5E-H Table). The mean time required for increasing echinocytosis was highly variable for Cy.003-(250) and Cy.007-(400) but this only reached significance for Cy.007-(400) (mean 98.2 s) which was more than double that of the control (mean 44.2 s)(S2E Fig and S5E Table). Similarly for the duration of maximum echinocytosis, Cy.007-(400) was highly variable but not significantly so compared to the control (S2F Fig and S5F Table). For the period of decreasing echinocyotsis, the mean duration for R5.008-(40), Cy.009-(200), Cy.003-(250) and Cy.007-(400) increased relative to the control but was only statistically significant for Cy.003-(250) (S2G Fig and S5G Table). Despite echinocytosis being a post invasion phenomenon, the antibodies that targeted PfRH5 and PfCyRPA were able to exert an inhibitory/delaying/slowing effect, particularly for Cy.007-(400). Echinocytosis is thought to be triggered by RBC membrane lipid perturbations caused during merozoite invasion: the invasion slowing effects of the antibodies may have increased these effects thereby extending the period of echinocytosis [20, 27].

### Overall order and morphology of normal early ring development

We next examined the potential effects of the PfRH5 and PfCyRPA antibodies on the differentiation of merozoites into ring-stage parasites. Current knowledge of new ring development is largely restricted to observations of merozoites spinning or oscillating immediately following invasion in *P. knowlesi* and *P. falciparum* [22, 27], with the most detailed description of the stages from an electron microscopy study on *P. knowlesi* [28]. Before examining the effects of PfRH5 and PfCyRPA antibodies we first performed live cell imaging on parasites treated with the control EBL 040-(400) and found that the intraerythrocytic merozoites begin to spin on average 16.4 s after the completion of invasion (Fig 1A). The spinning lasted an average of 116.3 s followed by the growth of a pseudopod like protrusion from the merozoite 159.2 s later. Almost a minute later (av. 57.2 s), the merozoite lost its rounded shape and became irregular and was defined as a “preliminary ring” (Fig 1A, S1 Table). Almost 10 minutes after this (av. 638 s) a fully amoeboid ring was formed, defined as the point at which all dense regions on the rings became fluid and mobile (Fig 1A, S1 Table).

In the presence of EBL 040-(400), early ring development occurred in nearly all merozoites which invaded and became intracellular, with 97.4% forming a pseudopod-like protrusion (Fig 4A, S4A Table). Of the original invaders, 88.3% continued differentiation into preliminary rings (Fig 4B, S4B Table) and 70% developed into amoeboid rings during the 20-minute observation period (Fig 4C, S4C Table). During this time, the parasites remained at the invasion site and were likely tethered to the erythrocyte surface with no occurrences of fission between the parasitophorous vacuole membrane and erythrocyte membrane.

**Fig 4.**
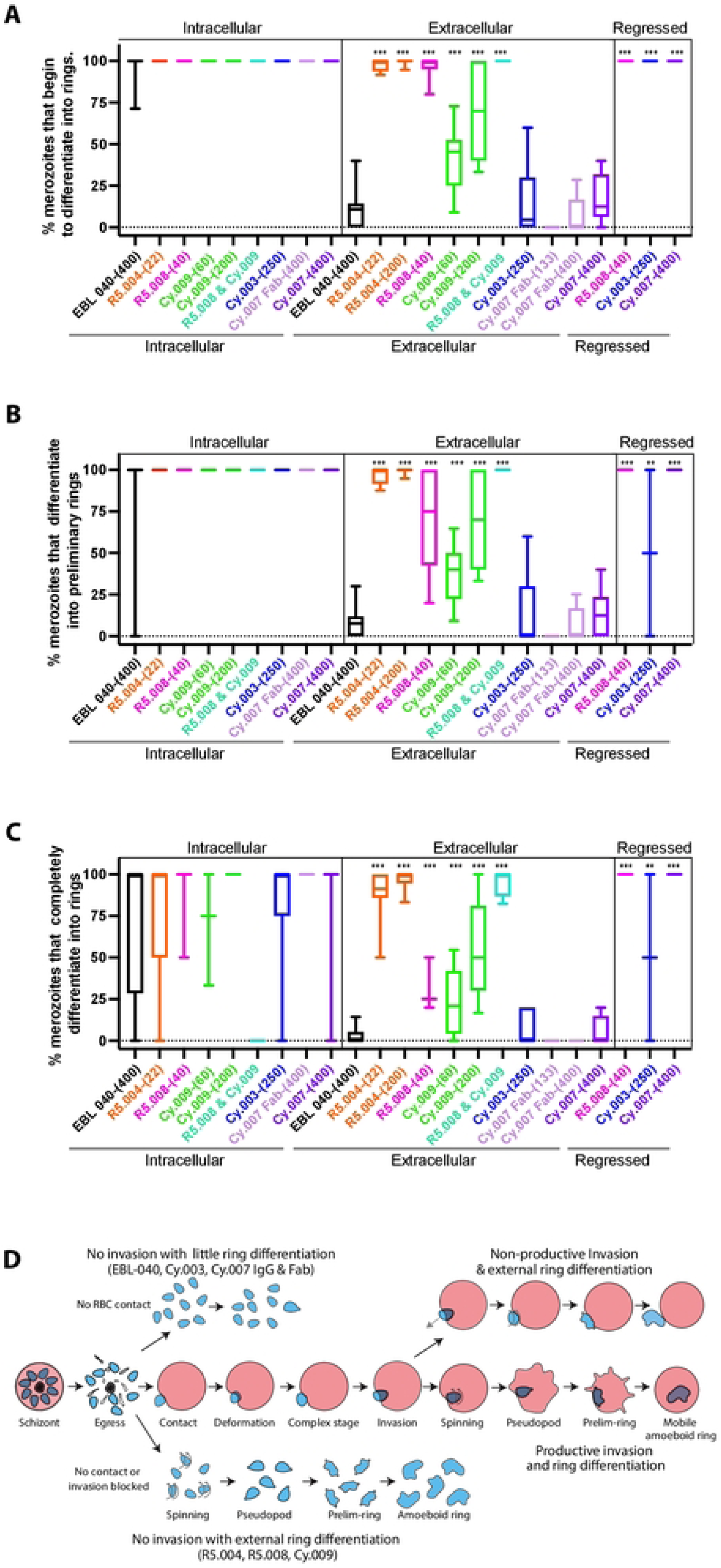
The effects of anti-PfRH5 and PfCyRPA antibodies upon the differentiation of intracellular and extracellular merozoites into ring-stage parasites. (A-C) The ability of the antibodies to stimulate or inhibit the differentiation of intracellular, extracellular and regressed merozoites into (A) early, (B) preliminary and (C) complete ring-stage parasites was assessed from observing live cell imaging videos of *Plasmodium falciparum* parasites. Full antibody names and concentrations (µg/mL) are indicated below bottom graph. The boxes indicate the median number of events and the 25% to 75% percentile, and whiskers show the range. Statistical analyses were performed using unpaired t tests in GraphPad PRISM V 9.0. The asterisks indicate where parasite mAbs have altered the number of events significantly from the EBL 040 intracellular or extracellular control (arrows) with *p<0.05, **p<0.01 and ***p<0.001. (D) Diagram summarizing the effects of the anti-PfRH5 and -PfCyRPA antibodies upon ring differentiation for intracellular and extracellular merozoites.

Most uninvaded, extracellular merozoites in the presence of EBL 040 IgG appeared unchanged over the 20-minute observation period. However, some extracellular merozoites began to spin with 10.7% of the original extracellular merozoites eventually developing pseudopods. Following this, 7.5% of the EBL 040 IgG treated merozoites lost their typically rounded shape and became more irregular. These so-called ‘preliminary rings’ then became fully amoeboid forms with 2.8% of the original EBL 040 IgG treated merozoites becoming amoebas (Fig 4A-C; S4A-C Table). As the pseudopods of the extracellular amoeba were not restrained within a parasitophorous vacuole, they became greatly elongated.

### Antibody mediated inactivation of extracellular merozoites

In the presence of anti-PfRH5 and -PfCyRPA mAbs, the percentage of merozoites beginning to differentiate into intracellular rings by developing pseudopods after invasion was very high and comparable to the control (Fig 4A, S4A Table). The percentages of merozoites that differentiated into intracellular preliminary rings and into complete rings in the presence of these mAbs was likewise similar to the EBL 040 control (Fig 4B and C and S4B and C Tables). A major difference was observed, however, for extracellular merozoites in R5.004-(22), R5.004-(200), R5.008-(40), Cy.009-(60) and Cy.009-(200), where most of the merozoites began to differentiate into rings by growing pseudopods compared to EBL 040, Cy.003-(250) and Cy.007 conditions (Fig 4A and D, S4A Tables, S1-S6 Videos). Extracellular merozoites developed pseudopods in 97.8% of cases in R5.004-(22), 99.5% in R5.004-(200), 96.7% in R5.008-(40), 71.1% in Cy.009-(200) and 100% of cases in the R5.008-(40)+Cy.009-(200) combination (Fig 4A, S4A Table, S2-S6 Videos). Following on, most of the PfRH5 and Cy.009 mAb treated extracellular merozoites developed into preliminary rings as they lost their rounded shape which reached 96.5% in R5.004-(22), 99.5% in R5.004-(200), 70.0% in R5.008-(40), 69.8% in Cy.009-(200) and 100% in the R5.008-(40)+Cy.009- (200) combination (Fig 4B, S4B Table, S2-S6 Videos).

The extracellular preliminary rings then differentiated into fully amoeboid rings in 89.0% of cases in R5.004-(22), 96.7% in R5.004-(200), 31.7% in R5.008-(40), 56.4% in Cy.009-(200) and 95.6% of cases in the R5.008-(40)+Cy.009-(200) combination (Fig 4C, S4C Table). The percentage of extracellular merozoites in R5.004-(22) and R5.008-(40), R5.004-(200) and the R5.008- (40)+Cy.009-(200) combination which develop into extracellular rings is comparable to invaded merozoites at every stage of ring development (Fig 4A-D, S4A-C Table).

In contrast, Cy.003, Cy.007 Fab, and Cy.007 IgG caused no additional extracellular ring development compared to the EBL 040 control (Fig 4A-D, S4A-C Tables, S7-S10 Videos). Cy.007 Fab-(400) was noteworthy as no extracellular merozoites reached the amoeboid ring stage (Fig 4C). While most conditions caused either nearly complete or close to no extracellular ring development, Cy.009-(60) caused intermediate levels of extracellular pseudopod (41.4%), preliminary ring (36.9%), and amoeboid ring (24.2%) development, leaving this group differing from the binary situation evident with other conditions (Fig 4A-D, S4A-C Tables).

### Antibody effects on regression

As mentioned previously, in the presence of R5.008-(40), Cy.003-(250), and Cy.007-(400), some merozoites regressed out of erythrocytes after invasion compared to no regression in EBL 040 (Fig 2D). Specifically, 22.78% of R5.008-(40) invasions regressed, 19.00% in Cy.003-(250) and 33.33% in Cy007-(400) (Figs 2D, S2D Table; S1, S3, S7 and S11 Video). We observed that most regressed merozoites became fully differentiated extracellular rings within the 20-minute observation (Fig 4A-C, S4A-C Table).

### Antibody effects on merozoite spinning

As indicated in Fig 1A, the first notable activity performed by merozoites after invasion is their spinning, oscillating, or twisting actions that have been implicated in helping sever the nascent PVM from the host cell plasma membrane [22, 27, 29]. To compare extracellular and intracellular ring development, we measured time from egress (when merozoite exposure to mAbs began) to the start of spinning (the first observable indication of ring conversion). On average, in the presence of EBL 040, intracellular merozoites began spinning 173.5 s after egress, more quickly than extracellular merozoites at 284.1 s (S3A Fig, S6A Table). Here, intracellular merozoites spun for nearly two minutes and extracellular merozoites for nearly 4 minutes (S3B, S6B Table). As a general observation, the mean times from egress to spinning for extracellular merozoites were shorter in the presence of the anti-PfRH5 mAbs than the extracellular control but this only reached significance for the R5.004 mAbs (Fig S3A, S6A Table). The times to spinning for the same extracellular anti-PfRH5 mAb treated merozoites were similar to the invaded intracellular EBL 040 treated merozoites (mean 173.5 s) with the exception of R5.004-(22) which could start spinning in as little as 21 s (mean 88 s, Fig S3A, S6A Table).

The duration of spinning in all extracellular merozoites developing into rings was comparable to that observed in intracellular EBL 040 except for parasites in R5.004-(200) and Cy.009-(200) which spun for less time and Cy.007-(400) where spinning lasted longer (S3B Fig, S6B Table). For extracellular merozoites in the presence of R5.004-(22), R5.004-(200) and Cy.009-(200), the duration of spinning was shorter than in the extracellular EBL 040 control (S3B Fig, S6B Table).

### Antibody effects on pseudopod formation

R5.004, R5.008-(40), and Cy.009 treated intracellular merozoites usually stopped spinning before the pseudopod became visible, similar to both EBL040 controls producing positive values (S3C Fig, S6C Table). In contrast, pseudopods were often visible before the end of spinning of intracellular merozoites in most Cy.003 and Cy.007 events, producing negative values (S3C Fig, S6C Table). For most of the antibody-treated extracellular merozoites, the first pseudopod was often visible before the end of spinning compared to intracellular merozoites. This was particularly so for extracellular merozoites treated with Cy.003 or Cy.007 suggesting merozoite development may be more dysregulated for extracellular compared to intracellular merozoites (S3C Fig).

### Antibody effects on preliminary ring formation

For intracellular merozoites, preliminary ring formation was defined as the time from the merozoite’s pseudopod first being visible until disruption of the rest of the merozoite’s plasma membrane (Fig 1A). For most intracellular merozoites in the presence of anti-PfRh5 and -PfCyRPA mAbs, the time to preliminary ring formation was comparable to the control antibody except for Cy.003 which took 4.5-fold longer (S3D Fig, S6D Table). For extracellular merozoites exposed to R5.004, R5.008 and Cy.009, it took less time for the merozoites to become preliminary rings than the extracellular EBA 040 control marked from when the first pseudopod was visible to when the merozoite became more irregularly shaped. In R5.004, R5.008 and Cy.009 IgGs, the extracellular merozoites became preliminary rings with similar times as the intracellular EBA 040 control (S3D Fig, S6D Table). For extracellular Cy.003 merozoites, preliminary development again took 4-fold longer than extracellular and intracellular controls (S3D Fig, S6D Table).

### Antibody effects on amoeboid ring formation

For PfRH5 and PfCyRPA antibody-treated intracellular parasites, the time from merozoite membrane disruption to complete ring formation was similar to the intracellular EBA 040 control (S3E Fig, S6E Table). Extracellular merozoites in the presence of R5.004-(200) took less time to develop into amoeboid rings than the extracellular control (S3E Fig). Our observations generally indicated that anti-PfRH5-exposed extracellular merozoites, which are efficiently neutralized and converted to ring-like forms, do so in similar stages and timings as merozoites which have successfully invaded. In contrast, the Cy.003 and Cy.007 treated merozoites both intracellular and extracellular tend to be more dysregulated often taking longer to complete the neutralization steps. Finally, in the presence of R5.008-(40), timing of regressed merozoite differentiation was comparable to that of successful invasions, while regressed Cy.003-(250) and Cy007-(400) merozoites had timing similar to that of extracellular merozoite differentiation (S3D and E Fig, S6D and E Table).

## Discussion

Live cell invasion imaging has previously played an important role in establishing how the invasion slowing anti-PfRH5 human mAb R5.011 could boost the potency of the invasion-neutralizing human mAb R5.016 [4]. Indeed, these data identified a mechanism of synergy between non-competing vaccine-induced antibody clones that bound different epitopes on PfRH5 [4]. We thus decided to apply this approach to study potential synergies between anti-PfRH5 and -PfCyRPA antibodies. We discovered that anti-PfRH5 mAbs generally appear to slow aspects of the pre-invasion complex phase during which the tight junction formed, while anti-PfCyRPA mAbs appear to slow the initial deformation stage and the downstream internalization stage where the merozoite penetrates its erythrocyte. An exception was Cy.009 which often behaved more like an anti-PfRH5 mAb rather than the other anti-PfCyRPA mAbs. Titration of R5.008 alone and in combination with Cy.009 indicated this anti-PfCyRPA mAb was able to synergistically boost the inhibitory capacity of R5.008 both in GIA assays and by live cell imaging. Hopefully these synergies can be clinically achieved considering recent experimental vaccinations of rats with PfRH5 and PfCyRPA antigens combinations did not produce IgGs that inhibited *in vitro* parasite growth much better than PfRH5 antigen antibody alone [30].

We also observed for the first time that PfRH5 and Cy.009 antibodies have effects on the post- invasion period (where newly invaded intraerythrocytic merozoites differentiate into ring-stage parasites). Unexpectedly, we found that the anti-PfRH5 mAbs and Cy.009 caused most of the merozoites that had not invaded to rapidly differentiate into amoeboid forms, which we believe are equivalent to ring-stage parasites. It is anticipated that the rapid induction of extracellular merozoites into rings could block the capacity of the parasites to attempt re-invasion and could increase the potency of antibodies that target PfRH5 and PfCyRPA.

One of the most inhibitory anti-PfRH5 human mAbs reported is R5.004 [4], which binds directly to the basigin binding site of PfRH5. R5.008 binds near the basigin binding site of PfRH5, probably sterically hindering PfRH5’s access to basigin. Using several measures, both PfRH5 mAbs reduced the number of invasions (per egress and invasions per merozoite-erythrocyte contact). This also applied to productive invasions where the merozoites did not regress during the observation period. For the limited number of invasions that were successful, our live cell imaging indicated both mAbs caused significant delay between the end of deformation and the start of internalization when PfRH5 is thought to bind basigin as part of the PCRCR complex [6, 8, 9, 20]. It therefore seems likely that during erythrocyte contact, there is competition between basigin and the anti-PfRH5 mAbs to gain access to PfRH5 with a threshold level of interaction between PfRH5 and basigin required for successful invasion. Competition between basigin and the anti- PfRH5 mAbs for PfRH5 likely delays time to reach the invasion threshold.

The anti-PfCyRPA mAbs were also very potent at inhibiting invasion using the invasions per egress and invasions per contact measurements with the Cy.007 Fabs being especially effective. This indicates that the Cy.007 Fab may be able to access its epitope far more effectively in the confined space between the merozoite apex and the RBC bound basigin complex [25] than the whole 150 kDa IgG even though the Fab would have less avidity than parental mAb. Of the invasions which did occur, Cy.009 behaved (in a dose-dependent manner) similarly to the anti- PfRH5 mAbs by delaying the start of invasion after the end of deformation, suggesting Cy.009 may inhibit PCRCR complex formation or the binding of PCRCR to basigin [4, 6]. Although Cy.009 does not occupy basigin’s binding site on PfRH5 it could sterically inhibit PfRH5 from binding to basigin, particularly in the crowded environment at the erythrocyte membrane where basigin binds RBC partner proteins PCMA or MCT1 [8, 25]. Cy.003 and Cy.007 on the other hand did not delay PCRCR complex formation or the binding of PCRCR to basigin but rather slowed down merozoite internalization. This could be due to the mAbs somehow slowing movement of the merozoite through the tight junction into the erythrocyte. The combination of the R5.008 and Cy.009 mAbs so greatly inhibited invasion that only one event was recorded. Here, Cy.009 appeared to function similarly to the potentiating mAb R5.011, in that it slowed the pre-invasion phase allowing more time for the neutralizing PfRH5 mAb to function [4]. Although Cy.009 did not appear to increase pre-invasion times as much as R5.011 [4], Cy.009 was much more inhibitory by itself and proved particularly effective in combination with R5.008. How mAb- induced interference of the PCRCR complex mechanistically blocks invasion will be explored later during the discussion of extracellular differentiation into rings.

Once merozoite internalization is complete, the erythrocyte starts to become an echinocyte about half a minute later. This is thought to be due to the deposition of lipids and other materials from the parasite rhoptries into the erythrocyte membrane causing asymmetry in the lipid bilayer, producing outward bending protrusions of the erythrocyte surface [20, 27]. Recently, high resolution lattice light sheet microscopy has revealed that echinocytosis probably begins much sooner after invasion than previously thought, evident as undulations of the erythrocyte membrane [27]. We concur that the time it takes for echinocytosis to reach its maximum and for the erythrocyte to return to its normal biconcave shape vary broadly [20, 27]. The only consistently observed effect of the antibodies upon echinocytosis was that it appeared to initiate more rapidly during treatment with the anti-PfRH5 and Cy.009 mAbs after invasion, possibly because invasion had been delayed.

The transformation of newly invaded merozoites into ring-stage parasites was described several decades ago for *P. knowlesi* with the characteristic steps of merozoite spinning, pseudopod growth and transformation of the circular shaped merozoite into an amoeboid ring [22, 28]. Here we examined the timing of these events in *P. falciparum* and found the conversion steps were conserved and that most invaded merozoites converted into rings. Spinning or oscillation of newly invaded *P. falciparum* merozoites have been previously noted and are thought to be a mechanical mechanism to promote severance of the parasitophorous vacuole membrane from the host cell membrane [27, 29]. Although we know little about merozoite spinning and the various downstream steps that result in conversion into rings, it is interesting that anti-PfRH5 and -PfCyRPA antibodies, particularly Cy.003 and Cy.007, could still influence ring differentiation (*e.g.,* increasing spinning duration) despite the merozoites having already invaded. Whether the antibodies ultimately reduce the successful transformation and growth of intraerythrocytic parasites is not yet known.

The most interesting phenomenon observed in the presence of the antibodies was the rapid transformation of extracellular merozoites into amoeboid forms we believe are equivalent to intraerythrocytic rings. This was most evident in the presence of anti-PfRH5 mAbs and Cy.009, where 50-100% of extracellular merozoites became rings compared to around 10% in the presence of the EBL 040 control IgG and the other anti-PfCyRPA mAbs. In the presence of anti- PfRH5 mAbs, Cy.009, and the R5.008 + Cy.009 combination, the time from merozoite egress and hence antibody exposure, to transformation into extracellular rings occurred with similar timing as successfully invaded merozoites. Extracellular ring development is particularly interesting because it would presumably inactivate and neutralize the extracellular merozoites, preventing them from attempting another round of invasion. As egressed merozoites have been estimated to have a half-life of 5 minutes, this could mean a substantial proportion of the extracellular merozoites could be inactivated while they are still invasion competent [31].

The mechanism by which the anti-PfRH5 and Cy.009 antibodies trigger ring development in extracellular merozoites is not yet understood but we believe that induction is potentially mimicking some naturally occurring step that takes place during normal invasion. It has been shown that the PCRCR complex and basigin are involved in triggering a calcium ion flux event at the apical end of merozoites that are about to invade erythrocytes [9, 20, 27]. This event is thought to be part of tight junction formation whereby the rhoptries and micronemes release proteins that form the ring-like junction with the erythrocyte surface through which the merozoite propels itself into the erythrocyte [9, 20, 32–34]. It is possible that binding of IgGs to PfRH5 and subsequent crosslinking mediated by the bivalent IgG molecules may mimic binding of PfRH5 with basigin at the erythrocyte surface and trigger a merozoite apical calcium ion flux that initiates a signaling cascade in the merozoite leading to ring development. Experiments with merozoites treated with fluorescent calcium dyes could help resolve if the PfRH5 mAbs are triggering apical calcium fluxes.

It is curious that the Cy.009 antibody also promotes extracellular merozoite ring development given the Cy.003 and Cy.007 are not any more effective than EBL 040 control. Cy.009 binds to the B1 and B2 propeller regions of PfCyRPA like the other PfCyRPA mAbs, so perhaps subtle differences in its angle of binding are responsible for Cy.009’s ring promoting activity [7]. It is therefore important to understand the fine specificity of the anti-PfCyRPA mAbs for their epitopes because even though they bind to same PfCyRPA blades, the epitopes do not overlap, and this could have important biological implications as observed here.

## Conclusions

Here we have demonstrated that antibodies to PfRH5 and PfCyRPA block merozoite invasion and that mAbs R5.008 and Cy.009, that are specific for each of these respective proteins, function synergistically. Live cell imaging in the presence of antibodies at concentrations that partly reduce invasion indicate that the anti-PfRH5 and Cy.009 mAbs increase the period of PfRH5- basigin invasion complex formation suggesting they might reduce the efficiency with which PfRH5 can functionally access basigin. The other anti-PfCyRPA mAbs increase the time taken for merozoites to internalize, suggesting they could sterically inhibit the speed with which the merozoite passes through the tight junction which could indicate the PCRCR complex still persists at the tight junction during invasion. The mAbs also influence speed and efficiency with which invaded merozoites can differentiate into intraerythrocytic ring-stage parasites which is surprising as the PCRCR complex functions well before this. The most unexpected finding was that anti-PfRH5 and Cy.009 mAbs trigger rapid differentiation of extracellular uninvaded merozoites into ring-like parasites. We next aim to determine if the differentiation of invasion competent merozoites into invasion-incapable extracellular rings is rapid enough and occurs at antibody concentrations low enough to majorly boost the protective immunity of PfRH5 and PfCyRPA antigen-based human vaccines and how this can be improved.

## Materials and Methods

### Growth Inhibition Activity Assays

Growth Inhibition activity (GIA) assays were performed to the protocol of the international GIA reference center at NIAID, NIH, USA [35]. Parasite cultures were synchronized using treatment with 5% sorbitol on the day before GIA assay set up. One-cycle GIA assays were performed at the indicated concentrations of mAbs. Biochemical measurement using a *P. falciparum* lactate dehydrogenase assay was used to quantify endpoint parasitemia which has been described previously [36] Percent GIA was calculated using the following equation where RBC are red blood cells:

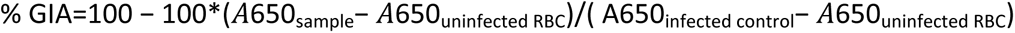

Assays were quality-controlled by inclusion of the anti-PfRH5 mAb 2AC7 as an in-plate control, and the anti-PfRH5 mAbs 2AC7, QA5, 9AD4 as external plate controls [37]. 5 mM EDTA is included as a positive control and a Zaire Ebolavirus glycoprotein-67 reactive IgG1 mAb (EBL 040) [21] was used as a negative isotype control for mAb samples.

To assess synergy/antagonism/additivity, two antibodies were used in combination. In all cases, one mAb was held at a constant concentration that on its own would be predicted to yield approximately 30% inhibition. A second mAb was then titrated alone to generate an inhibition curve. These data for the two mAbs (one titrated and one fixed concentration) are then used to calculate the predicted GIA of the combination by Bliss additivity [38]. This prediction was then compared to the real GIA result of the mAb mixture tested in parallel using the combination concentration of the two mAbs (*i.e.* the fixed + titrated amounts).

### Live cell imaging in the presence of antibodies which bind PfRH5 and PfCyRPA

Methods used are as described in [20], with the following exceptions: microscopy dishes used were 200 µL capacity, and the oxygen concentration used was 5% O_2_. Concentrations of antibody used were as follows: EBL 040 400 µg/mL, R5.004 22 µg/mL and 200 µg/mL, R5.008 40 µg/mL, Cy.009 60 µg/mL and 200 µg/mL, Cy.003 250 µg/mL, Cy.007 Fab fragment 133 µg/mL and 400 µg/mL, Cy.007 IgG 400 µg/mL.

R5.004 and R5.008 are anti-PfRH5 human IgG1 mAbs and have been reported previously [4]. Cy.009, Cy.007 and Cy.003 were provided by Icosagen AS through the Centre for AIDS Reagents repository at the National Institute for Biological Standards and Control, UK. These mAbs were produced through the European Commission FP7 EURIPRED project [39]. Cy.009, Cy.007 and Cy.003 were produced by Icosagen using HybriFree Technology. For clarity, these mAbs were renamed from their original EURIPRED consortium catalogue names, which should be cited for reagent requests (Cy.009 = 7B9#13; Cy.003 = 3B3#17; Cy.007 = 3A7#22).

### Statistical analysis

Data were analyzed using GraphPad Prism (GraphPad Software, version 9). For all unpaired t tests, two-tailed p values were considered significant if ≤0.05. Box and whisker plots indicate interquartile range with medians, and whiskers indicate minimum to maximum.

## Acknowledgements

G.E.W. was recipient of NHMRC Early Career Research Fellowship. We thank the Australian Red Cross Blood Bank for providing blood and the Victorian Operational Infrastructure Support Program received by the Burnet Institute. We are grateful for support from NHMRC grant 1092789. R.J.R. was funded by a Rhodes scholarship; the Wellcome Trust Infection, Immunology and Translational Medicine D. Phil programme; and a Canadian Institutes of Health Research Doctoral Foreign Study Award (FRN:157835). B.G.W. held a UK MRC PhD Studentships (MR/N013468/1). S.J.D. held a Wellcome Trust Senior Fellowship (106917/Z/15/Z). Cy.003, Cy.007, and Cy.009 were provided by Icosagen AS through the Centre for AIDS Reagents repository at the National Institute for Biological Standards and Control, UK. These mAbs were produced through the European Commission FP7 EURIPRED project (INFRA-2012-312661), funded by the European Union’s Seventh Framework Programme [FP7/2007—2013] under Grant Agreement No: 312661—European Research Infrastructures for Poverty Related Diseases (EURIPRED).

## Author contributions

G.E.W., roles: Conceptualization, Data Curation, Formal Analysis, Investigation, Methodology, Validation, Visualization, Writing – Original Draft Preparation, Writing – Review & Editing; R.J.R., roles: Investigation, Writing – Review & Editing; D.Q., roles: Investigation; A.M.L., roles: Investigation; M.G.D., role: Investigation, Writing – Review & Editing. C.B., role: Investigation, Writing – Review & Editing; B.G.W., role: Investigation, Methodology; B.S.C., role: Funding acquisition. S.J.D., Project Administration, Resources, Supervision, Writing – Review & Editing. P.R.G., roles: Conceptualization, Data curation, Funding acquisition, Investigation, Methodology, Project Administration, Resources, Supervision, Writing – Review & Editing

## Supplementary Figures

**S1 Fig.**
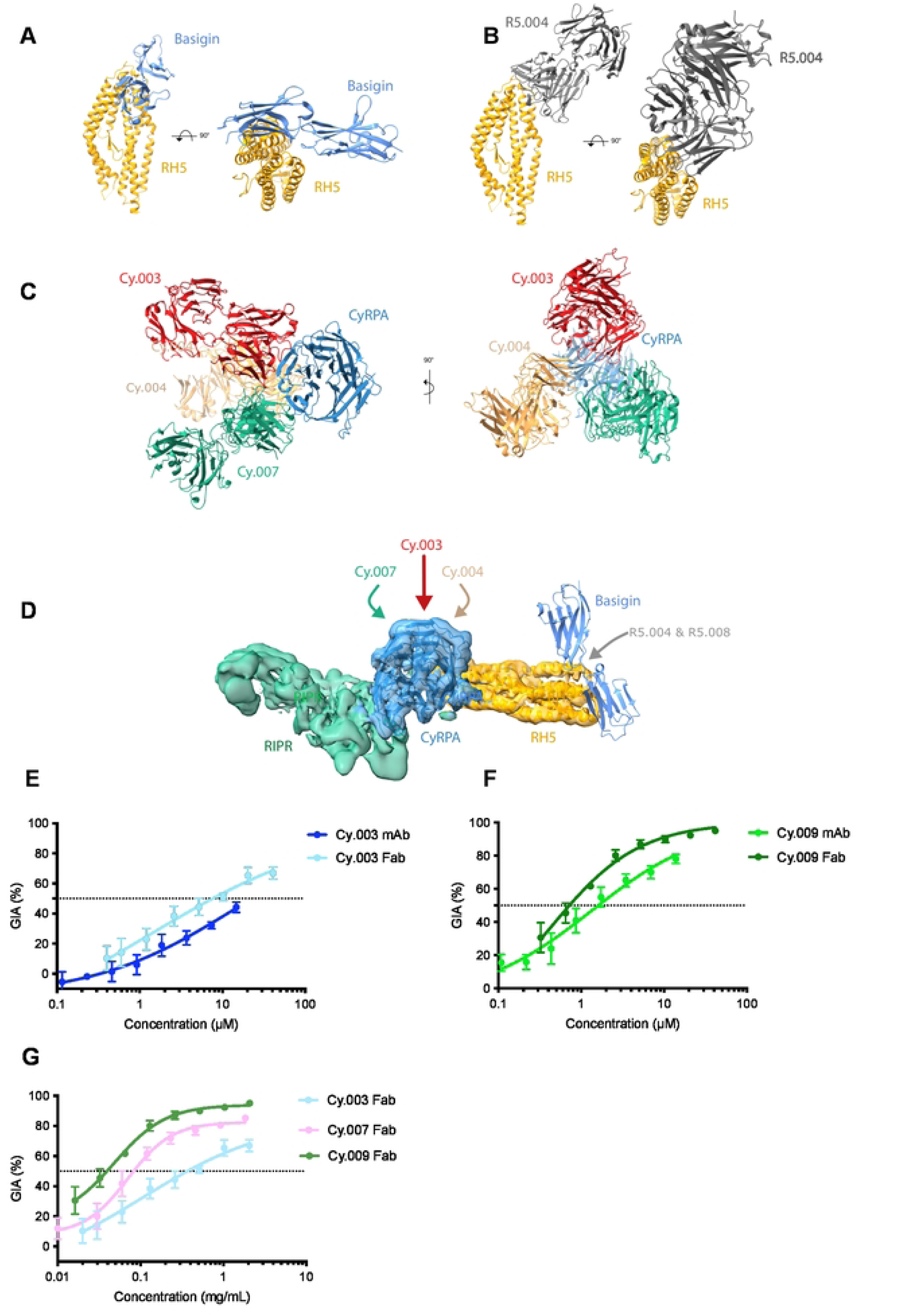
Epitopes recognized by anti-PfRH5 and -PfCyRPA Fabs and the growth inhibitory activity of PfCyRPA Fabs against the *Plasmodium falciparum* asexual blood stage. (**A**) Crystal structure of PfRH5 bound to basigin (protein data bank (PDB) ID: 4U0Q [40]). (B) Crystal structure of PfRH5 bound to R5.004 (PDB ID: 6RCU [4]). No structure of R5.008 is available, however both R5.004 and R5.008 compete with basigin for binding but do not compete with each other [4]. (C) Crystal structure of PfCyRPA bound to Cy.003, Cy.004, and Cy.007 (PDB ID: 7PI3 [19]). No structure is available for Cy.009 however Cy.004 and Cy.009 bind overlapping epitopes [19]. (D) Location of PfRH5 and PfCyRPA mAbs used in this study in the context of the PfRH5 (yellow), PfCyRPA (blue), PfRIPR (green) complex bound to basigin (composite image using PDB ID: 6MPV [8] & 4U0Q [40]). (E and F) Schizont stage *Plasmodium falciparum* 3D7 parasites were incubated with a micromolar dilution series of monoclonal antibodies (mAbs) and Fab fragments of Cy.003 and Cy.009 and grown for 40 h before parasite growth inhibitory activity (GIA) was quantified by measuring lactate dehydrogenase activity. (G) Comparison of GIA activities of anti-PfCyRPA Fabs.

**S2 Fig.**
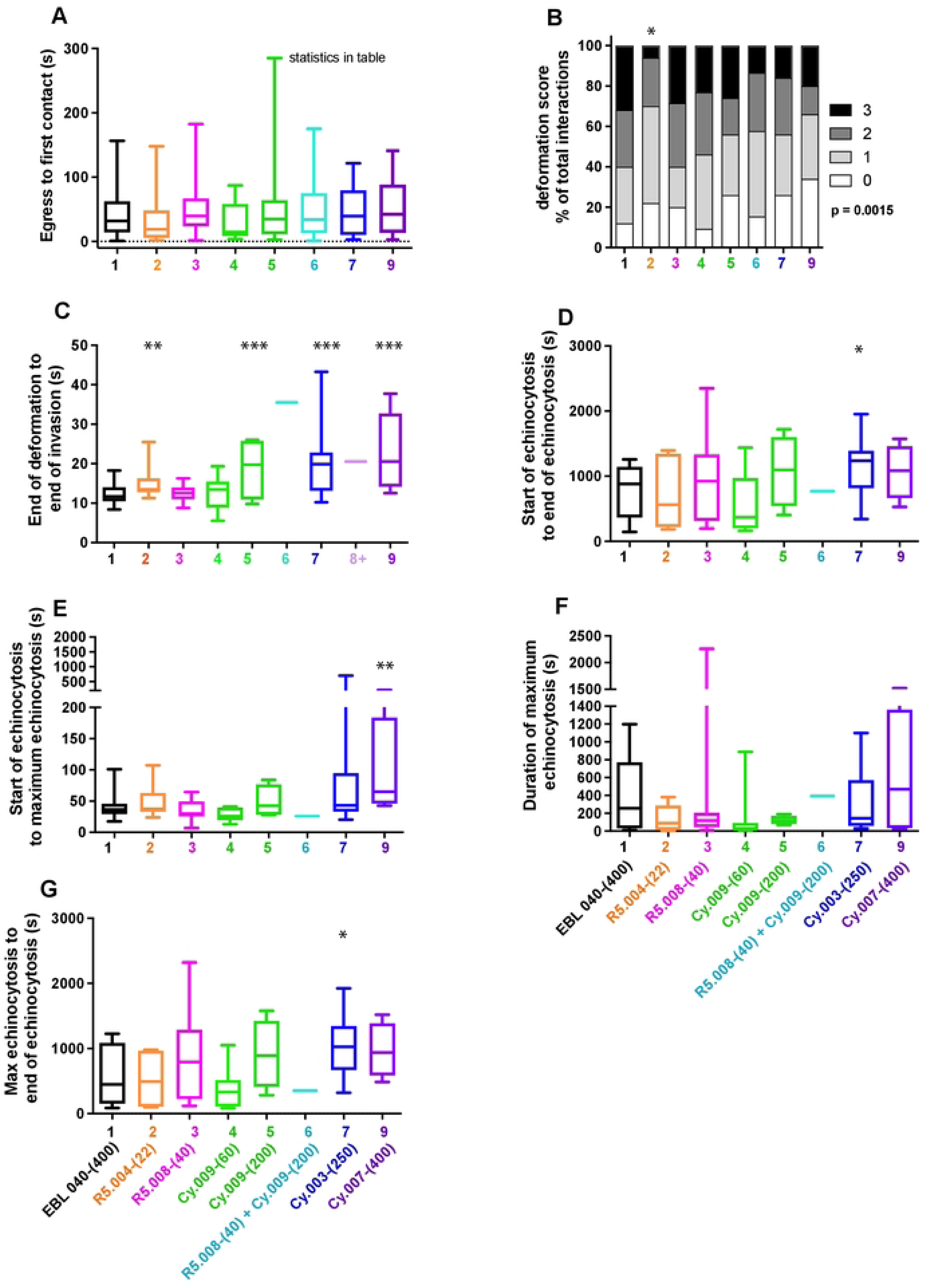
Quantification of the effects of antibodies to PfRH5 and PfCyRPA upon the invasion of erythrocytes by *Plasmodium falciparum* 3D7 merozoites. Video microscopy of several merozoite egress events was observed in the presence of antibodies with concentrations in µg/mL indicated in brackets. (A) The times from egress to first contact indicate were not significantly different indicating the imaging conditions were consistent. (B) The degree of deformation of merozoites on erythrocyte surfaces was quantified according to [20] in the presence of antibodies. R5.004-(22) caused significantly less deformation than the control or parasite antibodies using. (C-G) The timings of other invasion stage as indicated on the y-axes were measured using the antibody combinations names and concentrations (µg/mL) indicated below the x-axes. Antibody 8 is Cy.007 Fab-(133) and 8+ is Cy.007 Fab-(400). The boxes indicate the median number of events and the 25% to 75% percentile, and whiskers show the range. Statistical analyses were performed using unpaired t tests in GraphPad PRISM V 9.0. The asterisks indicate where parasite mAbs have altered the number of events significantly from the EBL 040 control with *p<0.05, **p<0.01 and ***p<0.001.

**S3 Fig.**
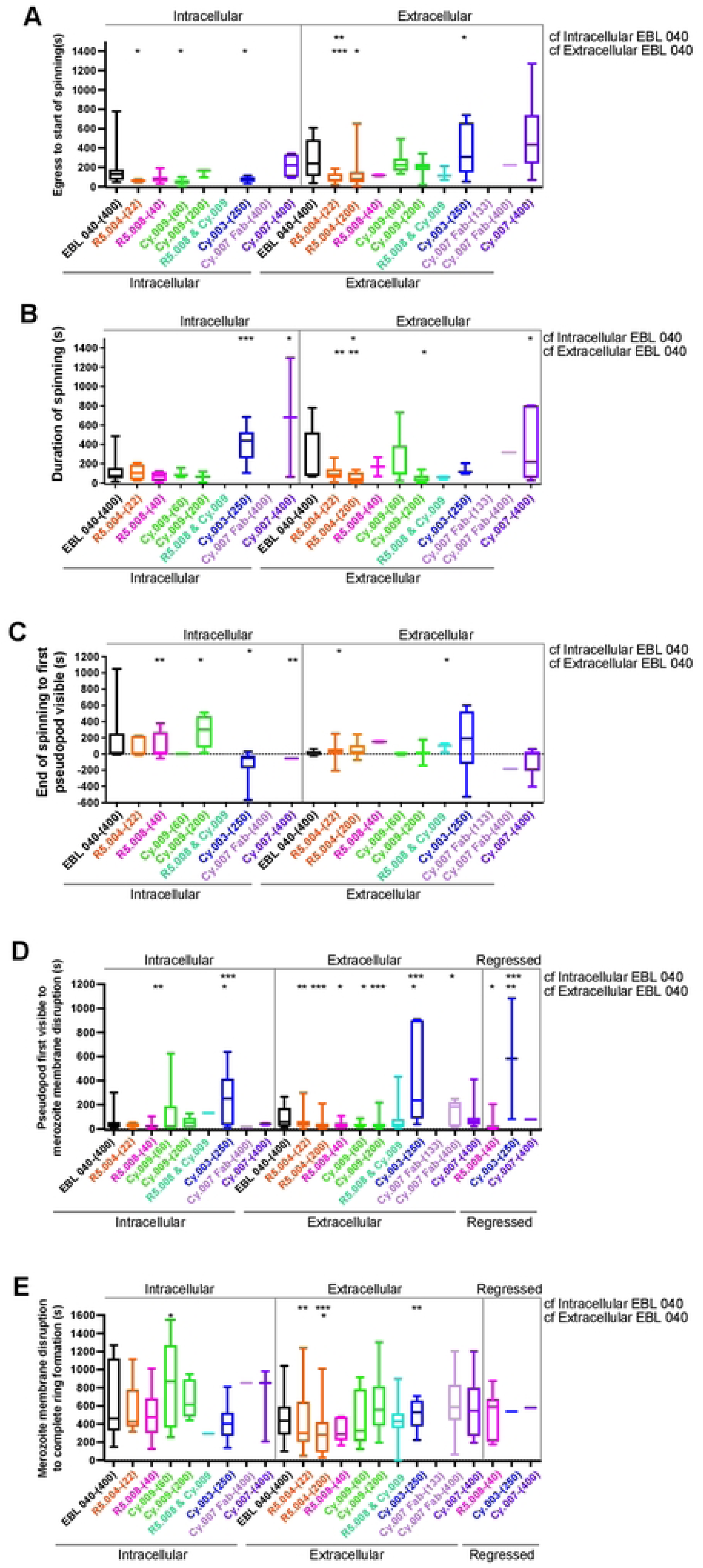
5 Anti-PfRH5 and -PfCyRPA antibodies influence the speed with which intracellular and extracellular merozoites differentiate into rings. (A) The speed with which merozoites started spinning after egress, (B) their duration of spinning, (C) from the end of spinning to pseudopod appearance, (D) pseudopod formation to disruption of merozoites surface until (E) complete ring formation respectively, were measured from live cell videos of cultured *Plasmodium falciparum*. Full antibody names and concentrations (µg/mL) are indicated below bottom graph. The boxes indicate the median number of events and the 25% to 75% percentile, and whiskers show the range. Statistical analyses were performed using unpaired t tests in GraphPad PRISM V 9.0. The asterisks indicate where parasite mAbs have altered the number of events significantly from the intracellular and extracellular EBL 040 controls with *p<0.05, **p<0.01 and ***p<0.001.

## Supplementary videos

**S1 Video EBL 040-(400). Live cell video of *Plasmodium falciparum* merozoites egressing from schizont in the presence of control EBL 040 IgG at 400 µg/mL.** First appearance of white arrow indicates merozoite invading erythrocyte and second appearance indicates differentiation of this merozoite into a ring stage form. Note, an invasion was chosen in which the invaded red blood cell did undergo echinocytosis so the merozoite’s development into a ring-stage parasite can be observed. Video playback is 10x live imaging speed.

**S2 Video R5.004-(22). Live cell video of *Plasmodium falciparum* merozoites egressing from schizont in the presence of R5.004 IgG at 22 µg/mL.** There are no erythrocyte invasions observed and black arrows indicates selected extracellular merozoites that are about to differentiate into amoeboid ring-like forms beginning about 3 minutes after egress. Video playback is 10x live imaging speed.

**S3 Video R5.008-(40). Live cell video of *Plasmodium falciparum* merozoites egressing from schizont in the presence of R5.008 IgG at 40 µg/mL.** Successful invasions are indicated with white arrows and black arrows indicate selected extracellular merozoites that begin to differentiate into amoeboid ring-like forms beginning about 5:40 minutes after egress. Video playback is 10x live imaging speed.

**S4 Video Cy.009-(60). Live cell video of *Plasmodium falciparum* merozoites egressing from schizont in the presence of Cy.009 IgG at 60 µg/mL.** No successful invasions were detected and the extracellular merozoites were observed to spin and begin to differentiate into amoeboid ring-like forms about 4 minutes after egress. Video playback is 10x live imaging speed.

**S5 Video Cy.009-(200). Live cell video of *Plasmodium falciparum* merozoites egressing from schizont in the presence of Cy.009 IgG at 200 µg/mL.** A successful invasion is indicated with a white arrow and note how long the invasion takes to commence after the merozoite makes first with its target erythrocyte. Black arrows indicate some of the extracellular merozoites that begin to differentiate into amoeboid ring-like forms about 5 minutes after egress. Video playback is 10x live imaging speed.

**S6 Video R5.008-(40) Cy.009-(200). Live cell video of *Plasmodium falciparum* merozoites egressing from schizont in the presence of R5.008 IgG at 40 µg/mL and Cy.009 IgG at 200 µg/mL.** White arrow indicates one successful invasion with an extended pre-invasion period. Black arrows indicate selected extracellular merozoites that begin to change into amoeboid ring-like forms at about 4 minutes post egress. Video playback is 10x live imaging speed.

**S7 Video Cy.003-(250). Live cell video of *Plasmodium falciparum* merozoites egressing from schizont in the presence of Cy.003 IgG at 250 µg/mL.** A successful merozoite invasion is indicated with a white arrow and at 17 s the time from the start to the end of invasion is longer than the control (10 s). Extracellular merozoites generally remained unchanged during the 10 minutes observation period. Video playback is 10x live imaging speed.

**S8 Video Cy.007 Fab-(133). Live cell video of *Plasmodium falciparum* merozoites egressing from schizont in the presence of Cy.007 Fab at 133 µg/mL.** No merozoite invasions were observed and most extracellular merozoites did not change during the 10 min imaging period. Video playback is 10x live imaging speed.

**S9 Video Cy.007 Fab-(400). Live cell video of *Plasmodium falciparum* merozoites egressing from schizont in the presence of Cy.007 Fab at 400 µg/mL.** No merozoite invasions were observed and most extracellular merozoites did not change during the 10 min imaging period. Video playback is 10x live imaging speed.

**S10 Video Cy.007-(400). Live cell video of *Plasmodium falciparum* merozoites egressing from schizont in the presence of Cy.007 IgG at 400 µg/mL.** No merozoite invasions were observed and most extracellular merozoites did not change during the 10 min imaging period. Video playback is 10x live imaging speed.

**S11 Video Cy.003-(250) Live cell video of *Plasmodium falciparum* merozoite regressing from invaded erythrocyte in the presence of Cy.003 IgG at 250 µg/mL.** Merozoite invading erythrocyte is indicated with a white arrow which changes to block to show the merozoite regressing from the erythrocyte. Video playback is 10x live imaging speed.

